# An fMRI dataset in response to “The Grand Budapest Hotel”, a socially-rich, naturalistic movie

**DOI:** 10.1101/2020.07.14.203257

**Authors:** Matteo Visconti di Oleggio Castello, Vassiki Chauhan, Guo Jiahui, M. Ida Gobbini

## Abstract

Naturalistic stimuli evoke strong, consistent, and information-rich patterns of brain activity, and engage large extents of the human brain. They allow researchers to compare highly similar brain responses across subjects, and to study how complex representations are encoded in brain activity. Here, we describe and share a dataset where 25 subjects watched part of the feature film “The Grand Budapest Hotel” by Wes Anderson. The movie has a large cast with many famous actors. Throughout the story, the camera shots highlight faces and expressions, which are fundamental to understand the complex narrative of the movie. This movie was chosen to sample brain activity specifically related to social interactions and face processing. This dataset provides researchers with fMRI data that can be used to explore social cognitive processes and face processing, adding to the existing neuroimaging datasets that sample brain activity with naturalistic movies.

## Background & Summary

In cognitive neuroscience the use of naturalistic stimuli such as commercial movies has advanced our understanding of the human brain. While simple, controlled experiments necessarily target a limited space of brain responses, naturalistic stimuli sample a much broader space ^1,2^. Naturalistic stimuli are better suited to engage participants and hold their attention ^3^. They evoke more reliable brain activity in comparison to controlled experiments where the same stimuli in the same conditions are repeated multiple times ^4–6^. Naturalistic stimuli allow researchers to compare highly similar brain responses across subjects ^1,4,5,7^, and to study how complex representations are encoded in brain activity ^2,8,9^.

Experiments with naturalistic stimuli are flexible because they can be analyzed with a variety of methods. Inter-Subject Correlation (ISC) ^10^ can be used to study the similarity of brain activity across subjects. Multivariate Pattern Analysis (MVPA) ^11,12^, including Representational Similarity Analysis (RSA) ^13,14^, can be used to investigate information in population responses embedded in patterns of brain activity. Voxelwise encoding models can be used with naturalistic stimuli to create predictive models of brain activity and quantify complex, multidimensional voxel tuning ^15–17^. Across subjects, brain activity is highly similar in response to naturalistic stimuli ^5^, and such brain responses can be used as a basis for functional alignment (e.g., Hyperalignment ^1,7,18^). Hyperalignment outperforms anatomical alignment for statistical analysis and, most importantly, preserves the information encoded in fine-grained topographies of brain activity, which facilitates the study of individual differences ^19–21^.

As an additional advantage, a single naturalistic fMRI dataset can be reused multiple times to answer different experimental questions with a variety of analytical methods. When datasets with responses to naturalistic stimuli are publicly shared, they can be used by many different laboratories and researchers to address their specific questions of interest (for example, see http://studyforrest.org/ ^22^). It is important, however, to use a variety of naturalistic stimuli to sample brain activity more broadly. For example, multiple naturalistic movies can be used to test whether the experimental results of interest generalize beyond a specific stimulus set.

For this reason, in this paper we describe and share a dataset where 25 subjects watched part of the feature film “The Grand Budapest Hotel” by Wes Anderson. Their brain activity was measured with a state-of-the-art 3T scanner (Siemens Prisma) at the Dartmouth Brain Imaging Center. This movie was chosen to sample brain activity specifically related to social interactions and face processing. The movie has a large cast with many famous actors. Throughout the story, the camera highlights many different faces and expressions, which are fundamental to understand the complex narrative of the movie. In a previous publication ^23^, part of this dataset has been used as a basis for hyperalignment to show that face-specific functional ROIs can be recovered in new subjects using existing data, paving the way for a novel method to recover functional ROIs in detail without using time-consuming localizer tasks. This dataset adds to the existing neuroimaging datasets that sampled brain activity with naturalistic movies ^22,24–26^ to provide researchers with fMRI data that can be used by those especially interested in social interactions and face perception.

## Methods

### Participants

Twenty-five participants (including three of the authors, 13 females, mean age 27.52 years ± 2.26 SD) took part in the experiment. All had normal or corrected-to-normal vision. Twenty-one participants used a custom-fitted CaseForge headcase (https://caseforge.co) to minimize head motion in the scanner (see Table 1). The study was approved by the Dartmouth Committee for the Protection of Human Subjects.

**Table 1.** Demographic information about subjects.

| Subject ID | Age | Sex | Headcase? |
| --- | --- | --- | --- |
| SID000005 | 29 | M | Yes |
| SID000007 | 25 | M | Yes |
| SID000009 | 28 | M | Yes |
| SID000010 | 29 | M | Yes |
| SID000013 | 26 | F | Yes |
| SID000020 | 28 | F | Yes |
| SID000021 | 31 | M | Yes |
| SID000024 | 31 | M | Yes |
| SID000025 | 29 | M | No |
| SID000029 | 25 | F | Yes |
| SID000030 | 29 | F | No |
| SID000034 | 25 | F | Yes |
| SID000050 | 27 | F | No |
| SID000052 | 24 | F | Yes |
| SID000055 | 30 | M | No |
| SID000114 | 28 | F | Yes |
| SID000120 | 25 | F | Yes |
| SID000134 | 29 | F | Yes |
| SID000142 | 26 | F | Yes |
| SID000278 | 29 | M | Yes |
| SID000416 | 28 | M | Yes |
| SID000499 | 29 | M | Yes |
| SID000522 | 27 | F | Yes |
| SID000535 | 22 | M | Yes |
| SID000560 | 29 | F | Yes |

### Stimuli

The full-length feature movie “The Grand Budapest Hotel” by Wes Anderson (DVD UPC 024543897385) was divided into six parts of different durations. The movie was split at scene cuts to maintain the narrative of the movie as intact as possible. The audio of the movie was post-processed using FFMPEG (https://www.ffmpeg.org) with an audio compressor filter to reduce the dynamic range and make dialogues clearer in the scanner. The code used to split and post-process the movie is available in the code repository.

### Procedure

Subjects took part in two experimental sessions, one behavioral and one in the fMRI scanner. In the behavioral session, participants watched the first part of the movie (approximately 46 minutes). Immediately after this session, participants went into the scanner and watched the remaining movie, divided into five parts. They were instructed to watch the movie without any additional task.

### Imaging session

The imaging session comprised one anatomical (T1w) scan, one gradient echo (GRE) fieldmap estimation scan, and five functional runs. During the anatomical scan participants watched the last five minutes of the first part of the movie—which they watched in the behavioral session—to calibrate the sound volume for the scanner. They were asked to use a button box to increase or decrease the volume so that they could easily hear the dialogue. The volume chosen by the subject was used throughout the session without further modifications. The functional runs had a different duration depending on the part of the movie and ranged from approximately 9 to 13 minutes. Each run was padded with a 10 s fixation period both at the beginning and the end of the run. In all but the first run, the movie started with at least 10 s that overlapped with the previous run. The movie was presented to the subjects on a back-projected screen, and subtended approximately 16.27 × 9.17 (W × H) degrees of visual angle. The audio was delivered to the subject through MR-compatible in-ear headphones (Sensimetrics model S14).

### Imaging parameters

All functional and structural volumes were acquired using a 3 T Siemens Magnetom Prisma MRI scanner (Siemens, Erlangen, Germany) with a 32-channel phased-array head coil at the Dartmouth Brain Imaging Center. Functional, blood oxygenation level-dependent (BOLD) images were acquired in an interleaved fashion using gradient-echo echo-planar imaging with pre-scan normalization, fat suppression, a multiband (i.e., simultaneous multi-slice; SMS) acceleration factor of 4 (using blipped CAIPIRINHA), and no in-plane acceleration (i.e., GRAPPA acceleration factor of 1): TR / TE = 1000 / 33 ms, flip angle = 59°, resolution = 2.5 mm^3^ isotropic voxels, matrix size = 96×96, FoV = 240×240 mm, 52 axial slices with full brain coverage and no gap, anterior–posterior phase encoding. At the beginning of each run, three dummy scans were acquired to allow for signal stabilization. At the beginning of the imaging session, a single dual-echo GRE (gradient echo) scan was acquired. This scan was used to obtain a fieldmap estimate for spatial distortion correction.

A T1-weighted structural scan was acquired using a high-resolution single-shot MPRAGE sequence with an in-plane acceleration factor of 2 using GRAPPA: TR / TE / TI = 2300 / 2.32 / 933 ms, flip angle = 8°, resolution = 0.9375 × 0.9375 × 0.9 mm voxels, matrix size = 256 × 256, FoV = 240 × 240 × 172.8 mm, 192 sagittal slices, ascending acquisition, anterior–posterior phase encoding, no fat suppression, 5 min 21 s total acquisition time.

### Preprocessing

The description of the anatomical and functional preprocessing (sections <u>Anatomical data preprocessing</u> and <u>Functional data preprocessing</u>) was automatically generated by fMRIprep ^27^, and it is copied here with minimal changes for style (see https://fmriprep.org/en/stable/citing.html#note-for-reviewers-and-editors for more information).

Results included in this manuscript come from preprocessing performed using *fMRIPrep* 20.1.1 ^27^ (RRID:SCR_016216), which is based on *Nipype* 1.5.0 ^28,29^ (RRID:SCR_002502).

#### Anatomical data preprocessing

The T1-weighted (T1w) image was corrected for intensity non-uniformity (INU) with *N4BiasFieldCorrection* ^30^, distributed with *ANTs* 2.2.0 ^31^ (RRID:SCR_004757), and used as T1w-reference throughout the workflow. The T1w-reference was then skull-stripped with a *Nipype* implementation of the *antsBrainExtraction.sh* workflow (from ANTs), using OASIS30ANTs as target template. Brain tissue segmentation of cerebrospinal fluid (CSF), white-matter (WM) and gray-matter (GM) was performed on the brain-extracted T1w using *fast ^32^* (FSL 5.0.9, RRID:SCR_002823).

Brain surfaces were reconstructed using *recon-all* ^33^ (FreeSurfer 6.0.1, RRID:SCR_001847) and the brain mask estimated previously was refined with a custom variation of the method to reconcile ANTs-derived and FreeSurfer-derived segmentations of the cortical gray-matter of Mindboggle ^34^ (RRID:SCR_002438). Volume-based spatial normalization to one standard space (MNI152NLin2009cAsym) was performed through nonlinear registration with *antsRegistration* (ANTs 2.2.0), using brain-extracted versions of both T1w reference and the T1w template. The following template was selected for spatial normalization: ICBM 152 Nonlinear Asymmetrical template version 2009c ^35^ (RRID:SCR_008796; TemplateFlow ID: *MNI152NLin2009cAsym*).

#### Functional data preprocessing

For each of the five BOLD runs per subject, the following preprocessing was performed. First, a reference volume and its skull-stripped version were generated using a custom methodology of *fMRIPrep*. A B0-nonuniformity map (or fieldmap) was estimated based on a phase-difference map calculated with a dual-echo GRE (gradient-recall echo) sequence, processed with a custom workflow of *SDCFlows* inspired by the *epidewarp.fsl* script (http://www.nmr.mgh.harvard.edu/~greve/fbirn/b0/epidewarp.fsl) and further improvements in HCP Pipelines ^36^. The fieldmap was then co-registered to the target EPI (echo-planar imaging) reference run and converted to a displacements field map (amenable to registration tools such as ANTs) with FSL’s *fugue* and other *SDCflows* tools. Based on the estimated susceptibility distortion, a corrected EPI (echo-planar imaging) reference was calculated for a more accurate co-registration with the anatomical reference. The BOLD reference was then co-registered to the T1w reference using *bbregister* (FreeSurfer) which implements boundary-based registration ^37^. Co-registration was configured with six degrees of freedom. Head-motion parameters with respect to the BOLD reference (transformation matrices, and six corresponding rotation and translation parameters) are estimated before any spatiotemporal filtering using *mcflirt* ^38^ (FSL 5.0.9). BOLD runs were slice-time corrected using *3dTshift* from AFNI 20160207 ^39^ (RRID:SCR_005927). The BOLD time-series were resampled onto the fsaverage surface (FreeSurfer reconstruction nomenclature).

The BOLD time-series (including slice-timing correction when applied) were resampled onto their original, native space by applying a single, composite transform to correct for head-motion and susceptibility distortions. These resampled BOLD time-series will be referred to as preprocessed BOLD in original space, or just preprocessed BOLD.

Several confounding time-series were calculated based on the preprocessed BOLD: framewise displacement (FD), DVARS and three region-wise global signals. FD and DVARS are calculated for each functional run, both using their implementations in *Nipype* (following the definitions by Power et al. ^40^). The three global signals are extracted within the CSF, the WM, and the whole-brain masks. Additionally, a set of physiological regressors were extracted to allow for component-based noise correction (*CompCor* ^41^). Principal components are estimated after high-pass filtering the preprocessed BOLD time-series (using a discrete cosine filter with 128s cut-off) for the two *CompCor* variants: temporal (tCompCor) and anatomical (aCompCor). tCompCor components are then calculated from the top 5% variable voxels within a mask covering the subcortical regions. This subcortical mask is obtained by heavily eroding the brain mask, which ensures it does not include cortical GM regions. For aCompCor, components are calculated within the intersection of the aforementioned mask and the union of CSF and WM masks calculated in T1w space, after their projection to the native space of each functional run (using the inverse BOLD-to-T1w transformation). Components are also calculated separately within the WM and CSF masks. For each CompCor decomposition, the *k* components with the largest singular values are retained, such that the retained components’ time series are sufficient to explain 50 percent of variance across the nuisance mask (CSF, WM, combined, or temporal). The remaining components are dropped from consideration.

The head-motion estimates calculated in the correction step were also placed within the corresponding confounds file. The confound time series derived from head motion estimates and global signals were expanded with the inclusion of temporal derivatives and quadratic terms for each ^42^. Frames that exceeded a threshold of 0.5 mm FD or 1.5 standardized DVARS were annotated as motion outliers. All resamplings can be performed with a single interpolation step by composing all the pertinent transformations (i.e. head-motion transform matrices, susceptibility distortion correction when available, and co-registrations to anatomical and output spaces). Gridded (volumetric) resamplings were performed using *antsApplyTransforms* (ANTs), configured with Lanczos interpolation to minimize the smoothing effects of other kernels ^43^. Non-gridded (surface) resamplings were performed using *mri_vol2surf* (FreeSurfer).

Many internal operations of *fMRIPrep* use *Nilearn* 0.6.2 ^44^ (RRID:SCR_001362), mostly within the functional processing workflow. For more details of the pipeline, see the section corresponding to workflows in *fMRIPrep*’s documentation (https://fmriprep.readthedocs.io/en/latest/workflows.html).

#### Functional data denoising

The functional data preprocessed by fMRIprep was then denoised using custom Python scripts. The following nuisance parameters were regressed out from the functional time series using ordinary least-squares regression: six motion parameters and their derivatives, global signal, framewise displacement ^40^, the first six noise components estimated by aCompCor ^41^, and polynomial trends up to second order. All metrics of interest were computed on data denoised as described, either in volume space or in surface space. No additional spatial smoothing or temporal filtering was performed.

### Hyperalignment

We functionally aligned the functional data using whole-brain searchlight hyperalignment ^1,7,18,20^. The functional data projected to the fsaverage surface template and resampled to a low-resolution surface (10,242 vertices per hemisphere, approximately 3 mm resolution) was split in two separate datasets to perform hyperalignment and compute quality metrics on independent splits. The first split included runs 1-3, and the second split included runs 4 and 5. Transformation matrices were determined for disc searchlights of radius 15 mm, ignoring vertices in the medial wall. One subject (*sub-sid000009*) was used as the reference subject to create the hyperalignment common space. Data was z-scored before and after hyperalignment to normalize variance.

### Estimation of temporal signal-to-noise ratio (tSNR)

We first computed tSNR for each preprocessed functional run using data in each subject’s anatomy without template normalization, which would smooth the data spatially and affect tSNR. For each voxel, tSNR was calculated within each separate run as the temporal mean divided by the temporal standard deviation ^45^. A tSNR map was generated for each subject by computing the median tSNR across runs within each voxel. To qualitatively visualize how tSNR varied according to brain areas and generate a group tSNR map, the same analysis was performed with functional data resampled to the fsaverage surface.

### Inter-Subject Correlation

Inter-Subject Correlation was computed to estimate what proportion of the brain signal in response to the movie was consistent across subjects ^10^. The BOLD time series were projected to the template surface fsaverage, so that the data were spatially matched across subjects. Each subject’s data in a cortical node was correlated to the average time-series of the other 24 subjects in the same cortical node. This generated a map that quantifies the similarity of an individual subject’s response with the group response. The procedure was repeated for all subjects, and a median ISC map was computed at the group level (see Figure 3).

### Time-segment classification

Time-segment classification was used to estimate how much signal is available in local patterns of brain activity across subjects. First, functional data projected to the fsaverage template was hyperaligned (see Methods) with *sub-sid000009* as the reference subject. We used a nearest-neighbor classifier to distinguish between 15 s segments of the movie across subjects (chance level < 0.1 %). The movie segments were 1 TR apart and could have overlaps (see previous publications for more details ^18,19^). Classification was performed within surface searchlights with a radius of 10 mm. The data from 24 subjects was averaged and used as a training set, and the classifier was tested on the left-out subject. This process was repeated for all 25 subjects, and a final map was created by averaging across the 25 cross-validation folds.

## Data Records

The raw data was standardized following the Brain Imaging Data Structure ^46^ (version 1.3.0) to facilitate data sharing and the use of tools such as fMRIprep and MRIQC ^47^. The dataset is available on OpenNeuro (https://openneuro.org/datasets/ds003017), and can be easily downloaded using DataLad ^48^ from http://datasets.datalad.org/?dir=/labs/gobbini. While we cannot share the raw stimuli for copyright reasons, we provide the scripts that were used to preprocess the stimuli with all the information needed for other researchers to generate the same stimuli. We also share presentation, preprocessing, and analyses scripts in the github repository (https://doi.org/10.5281/zenodo.3942173, https://github.com/mvdoc/budapest-fmri-data).

## Technical Validation

The dataset was validated using different metrics that quantify data quality in separate domains. We analyzed the amount of subjects’ motion to quantify potential noise in the data caused by subjects’ behavior. We estimated tSNR for each voxel separately to make sure that all subjects had comparable levels of SNR and to highlight areas with low SNR. We computed Inter-Subject Correlation (ISC) as a metric that is specific to experiments with naturalistic paradigms. We consider ISC as a sanity check that the stimulus generated similar brain responses across subjects. All the metrics described so far provide information about data quality at the level of single voxels or surface nodes. To quantify data quality for multivariate analyses, we functionally aligned the data using searchlight hyperalignment and performed time-segment classification across subjects.

We first quantified motion in the dataset by inspecting the motion parameters estimated by fMRIprep (see Methods). Overall subject’s motion was low. The median framewise displacement across subjects was 0.09 mm (minimum median across subjects of 0.06 mm, max 0.19 mm, see Figure 1). Across subjects, the median percentage of volumes marked as motion outliers by fMRIprep was 2.72% (min 0.03 %, max 22.72 %), with 20 subjects out of 25 having less than 5% volumes marked as outliers (fMRIprep defines an outlier as a volume in which framewise displacement is greater than 0.5 mm or standardized DVARS is greater than 1.5; see Methods.)

**Figure 1.**
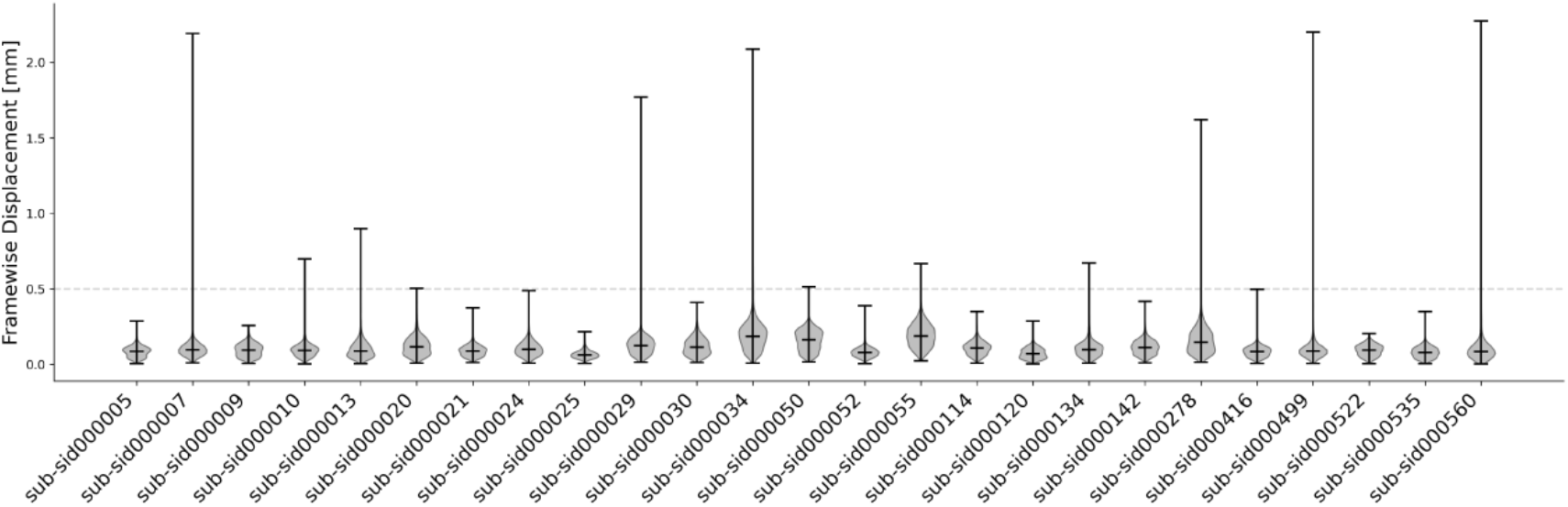
Framewise displacement for each subject across all runs. Subject motion was low in this dataset, as indicated by a median framewise displacement well below 0.5 mm for all subjects (the median value across subjects was 0.09 mm, minimum median across subjects of 0.06 mm, max 0.19 mm). Twenty out of 25 subjects had less than 5% volumes marked as motion outliers (fMRIprep defines an outlier as a volume in which framewise displacement is greater than 0.5 mm or standardized DVARS is greater than 1.5; see Methods.)

We estimated temporal SNR for all subjects, both in the subject’s own anatomical space (to reduce interpolations that can affect tSNR) and in the fsaverage template space for a qualitative assessment of tSNR across cortical areas. Temporal SNR is expected to vary across areas due to signal susceptibility artifacts, differences in anatomy across subjects, and overall subject arousal levels during the scan ^45^. The mean whole-brain tSNR across subjects was 74.42 ± 3.91, which is comparable to previous datasets ^49,50^. As expected, temporal SNR varied across areas, with higher tSNR in dorsal areas, and lower tSNR in anterior temporal cortex and orbito-frontal cortex (see Figure 2).

**Figure 2.**
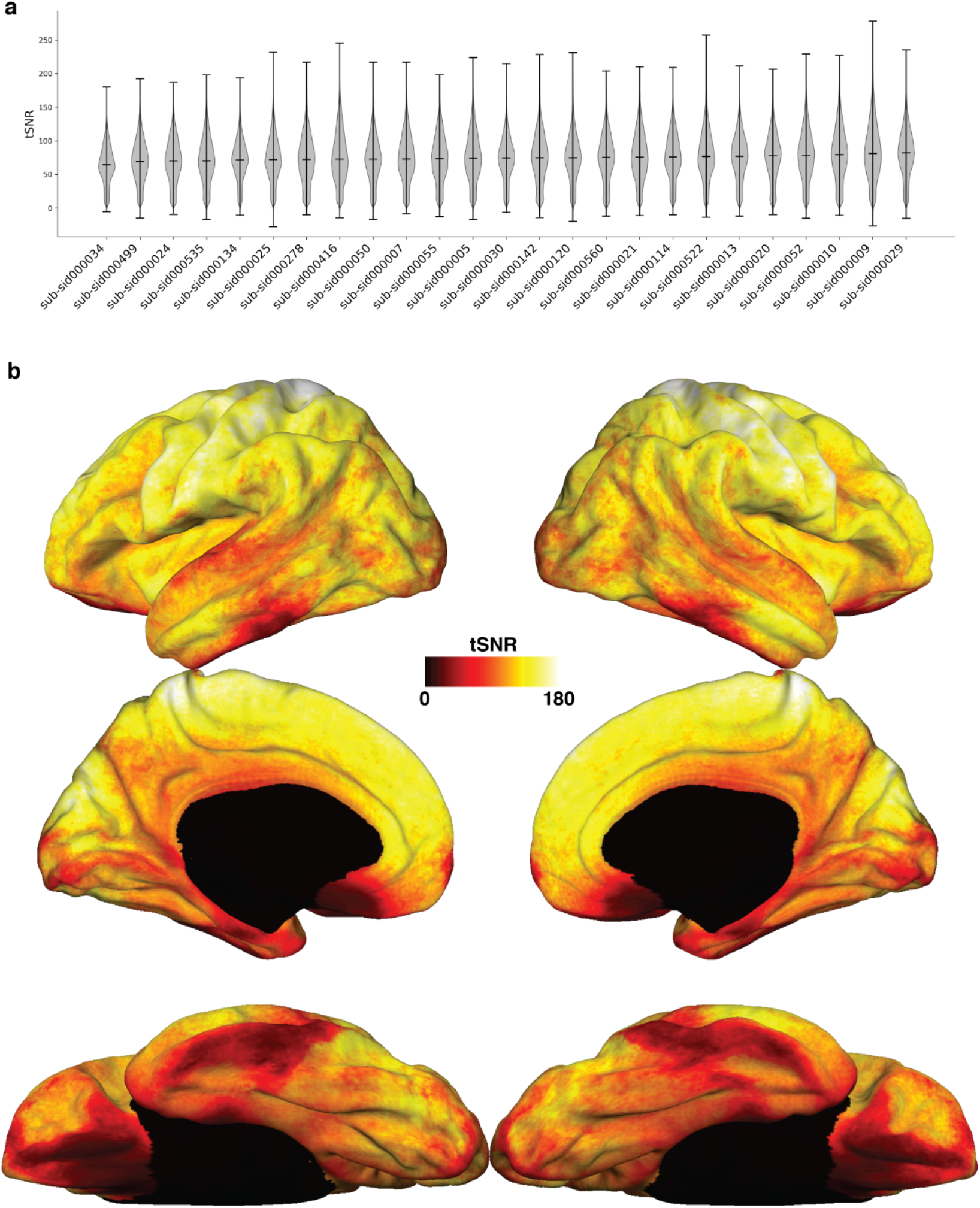
Temporal SNR across subjects. a) Violin plots showing tSNR values across the brain. For each subject, a tSNR map was first generated by computing the median tSNR value across runs within each voxel. This plot shows the distribution of values in the tSNR map within a brainmask, computed in each subject’s volumetric anatomical space. Subjects are ordered in increasing median tSNR. Across subjects, the mean tSNR was 74.42 ± 3.91. b) Median tSNR across subjects computed on data that was projected to the template surface fsaverage. As expected, areas closer to air-tissue boundaries such as the anterior temporal lobe and orbito-frontal cortex show signal dropout, while tSNR is high across the whole cortex.

We used Inter-Subject Correlation to highlight areas where brain activity in response to the movie was similar across subjects. As expected from an audio-visual movie, visual and auditory areas showed the highest ISC values (see Figure 3). In addition, areas known to process social information such as precuneus, temporo-parietal junction (TPJ), and medial prefrontal cortex (MPFC) ^51^ also showed positive ISC values. We speculate that this can reflect processing of the rich social information present in the movie, but future analyses might be required to investigate what further representations are encoded in these brain areas. Note that the ISC results in Figure 3 provide a lower bound of what can be obtained with this dataset. The analysis that we report was performed after anatomical alignment, which is known to be suboptimal for between-subject analyses such as ISC when compared to hyperalignment ^1^.

**Figure 3.**
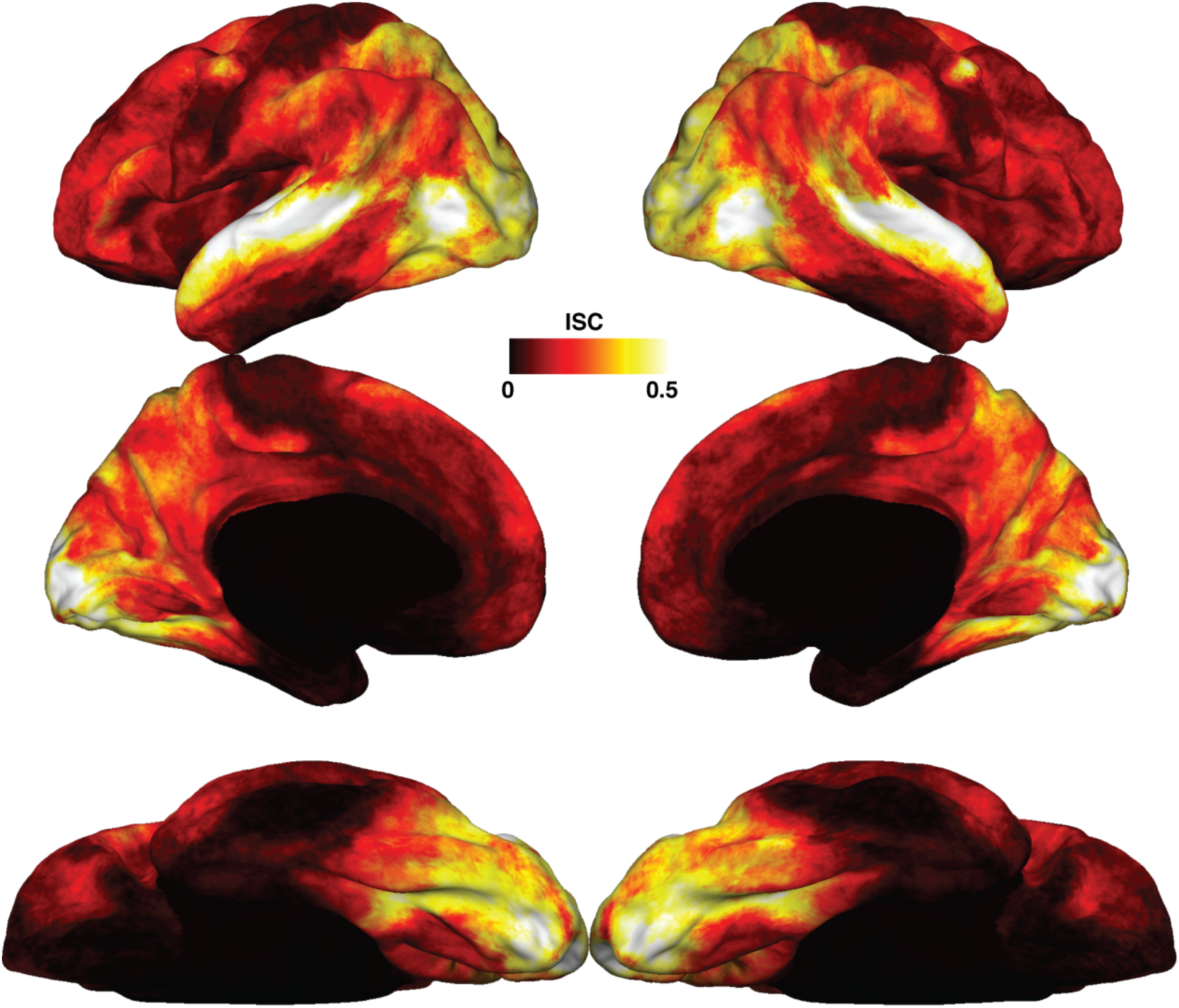
Inter-subject functional correlation. As expected from an audio-visual movie, visual and auditory areas showed the largest correlation in brain responses across subjects. Additional areas belonging to the theory-of-mind network, such as precuneus, temporo-parietal junction (TPJ) and medial prefrontal cortex (MPFC) also showed high correlation across subjects, as well as prefrontal areas, possibly highlighting the richness in social information available in the movie used for this dataset.

Finally, we performed between-subject time-segment classification to highlight areas whose patterns encode shared information across subjects. We first split the movie data into two independent sets (split 1: runs 1-3; split 2: run 4, 5). Then, we used data from one split to functionally align the subject’s data with whole-brain surface-searchlight hyperalignment ^1,7,18,20^. The data from the other split was then used to classify 15 s time-segments across participants. The process was repeated for both splits (see Figure 4). The average searchlight classification accuracy was 16.64%, and the maximum accuracy was 71.24%, with chance level less than 0.1 % (split 1: mean accuracy 18.91%, max accuracy 77.43%; split 2: mean 14.37%, max 65.05%). Classification accuracy was higher than chance level across the whole cortex. The highest accuracy values were found in visual and auditory cortex, but also in prefrontal and medial areas such as precuneus and medial prefrontal cortex.

**Figure 4.**
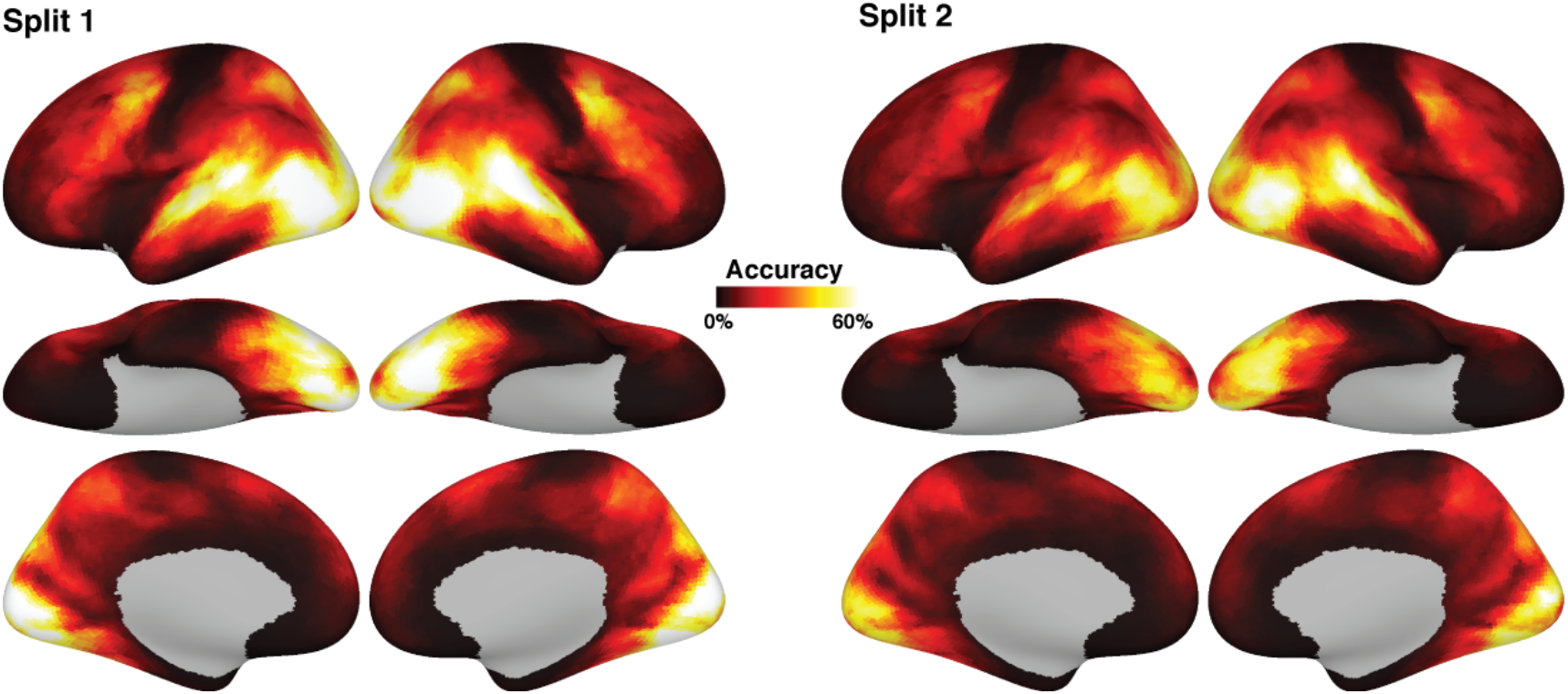
Between-subject time-segment classification on hyperaligned data. The left panel (split 1) shows results obtained from hyperaligning on the first half of the data (runs 1-3), and classifying on the second half (runs 4, 5). The right panel shows the complementary analysis, that is, hyperaligning on the second half of the data, and classifying on the first half. Despite differences in absolute classification values due to differences in amount of data, the results are qualitatively similar. The highest classification values could be found in visual and auditory areas, as well as theory-of-mind areas such as precuneus, TPJ, and MPFC, and also prefrontal areas.

These analyses validate the quality of this dataset for both univariate and multivariate analyses. We found evidence of overall good subject compliance, as reflected by low motion during scanning, as well as comparable tSNR levels across subjects. Inter-Subject correlation analysis and time-segment classification analyses both revealed shared information in visual and auditory areas, as well as the default mode network ^52,53^, which also plays a role in theory-of-mind processes ^51,54,55^.

## Code Availability

All code is available in the github repository ^56^ https://github.com/mvdoc/budapest-fmri-data. The code includes scripts to process the stimuli, presentation scripts, and scripts for the analyses presented in this paper. The scripts rely heavily on open source Python packages such as PyMVPA ^57^, nilearn ^44^, pycortex ^58^, scipy ^59^, and numpy ^60^.

## Acknowledgements

This project was supported by the NSF award #1835200 to M. Ida Gobbini. We would like to thank Jim Haxby, Yaroslav Halchenko, Sam Nastase, and the members of the Gobbini and Haxby lab for helpful discussions during the development of this project.

## Author contributions

MVdOC designed the experiment, wrote the presentation code, collected and analyzed the data, and wrote the manuscript.

VC collected and analyzed the data, and provided critical input to the manuscript.

GJ analyzed the data and provided critical input to the manuscript.

MIG designed the experiment, obtained funding, and edited the manuscript.

All the authors read and approved the manuscript.

